# CD147 (*BSG*) but not *ACE2* expression is detectable in vascular endothelial cells within single cell RNA sequencing datasets derived from multiple tissues in healthy individuals

**DOI:** 10.1101/2020.05.29.123513

**Authors:** C Ganier, X Du-Harpur, N Harun, B Wan, C Arthurs, NM Luscombe, FM Watt, MD Lynch

## Abstract

Coronavirus disease 2019 (COVID-19) is caused by severe acute respiratory syndrome coronavirus 2 (SARS-CoV-2) and is associated with a wide range of systemic manifestations. Several observations support a role for vascular endothelial dysfunction in the pathogenesis including an increased incidence of thrombotic events and coagulopathy and the presence of vascular risk factors as an independent predictor of poor prognosis. It has recently been reported that endothelitis is associated with viral inclusion bodies suggesting a direct role for SARS-CoV-2 in the pathogenesis. The ACE2 receptor has been shown to mediate SARS-CoV-2 uptake and it has been proposed that CD147 (BSG) can function as an alternative cell surface receptor. To define the endothelial cell populations that are susceptible to infection with SARS-CoV-2, we investigated the expression of ACE2 as well as other genes implicated in the cellular entry of SARS-Cov-2 in the vascular endothelium through the analysis of single cell sequencing data derived from multiple human tissues (skin, liver, kidney, lung and intestine). We found that CD147 (*BSG*) but not *ACE2* is detectable in vascular endothelial cells within single cell sequencing datasets derived from multiple tissues in healthy individuals. This implies that either *ACE2* is not expressed in healthy tissue but is instead induced in response to SARS-Cov2 or that SARS-Cov2 enters endothelial cells via an alternative receptor such as CD147.

## INTRODUCTION

We read with interest the report of Varga et al (Varga et al. 2020) showing endothelitis in association with viral inclusion bodies in endothelium from multiple organs of patients with coronavirus disease 2019 (COVID-19). This finding is of great relevance in view of multiple lines of evidence supporting a role for vascular endothelial dysfunction in the pathogenesis of COVID-19 including the presence of cardiovascular risk factors as an independent predictor of severe disease (Zhou et al. 2020), the high incidence of thrombotic complications (Klok et al. 2020) and the presence of coagulopathy (Tang et al. 2020). An understanding of the mechanism of endothelitis may shed light on the diverse systemic manifestations of COVID-19 and suggest potential therapeutic approaches.

The ACE2 receptor has been shown to mediate uptake of the virus responsible for COVID-19, severe acute respiratory syndrome coronavirus 2 (SARS-CoV-2), in human cells (Hoffmann, Kleine-Weber, Schroeder, et al. 2020) and previous reports have suggested expression of ACE2 in vascular endothelial cells (Lely et al. 2004; Hamming et al. 2004) and heart (Ferrario et al. 2005). In order to define the endothelial cell populations that are susceptible to infection with SARS-CoV-2, we investigated the expression of ACE2 in the vascular endothelium through the analysis of single cell sequencing data derived from multiple human tissues (skin, liver, kidney, lung and intestine).

## METHODS

Human lung and liver data were derived from the Human Cell Atlas project (Regev et al. 2018). Other data was publicly available (Liao et al. 2020; Aizarani et al. 2019; Travaglini et al. 2020; Y. Wang et al. 2020; Tabib et al. 2018). Single cell sequencing data is shown as dot plots and UMAP plots - in the latter each dot represents an individual cell and cells with similar expression patterns cluster together. Cell types were annotated according to markers as defined within the original publications. Vascular endothelium is identified by established markers including PECAM1, FLT1, VWF, TIE1, KDR, CD34 and CLDN5 (Leeuwenberg et al. 1992; Lee et al. 2015). We could identify endothelial cells in the lung, liver and skin but were unable to unambiguously identify endothelial cells in the intestine and kidney datasets.

## RESULTS

With the exception of enterocytes in the colon, the level of *ACE2* expression was very low across all of these datasets (Figure 1A, C and Table 1) and was not detected at a significant level in endothelial cells (Figure 1A, B, C and Table 1). In order not to inadvertently exclude any *ACE2*-expressing cells, we show plots without the usual filtering employed for single cell sequencing data (Supplementary methods). We also examined expression of *TMPRSS2,* a serine protease that is required for the priming of the spike protein of SARS-CoV-2 (Hoffmann, Kleine-Weber, Schroeder, et al. 2020). This was not expressed at a high level in endothelial cells in any of the tissues that we analysed (Figure 1A, D and Table 1). However it was detected in epithelial cells within the lung, liver and colon (Supplementary Figure 1 and Table 1). *TMPRSS2* was also expressed in hepatocytes. *CTSL* and *CTSB* encode Capthesin L and B, respectively and are alternative proteases which can mediate the priming of the spike protein (Hoffmann, Kleine-Weber, Krüger, et al. 2020). These are expressed across a range of cell types including endothelial cells in these different tissue types (Table 1).

**Table 1.** Expression of *ACE2, BSG* and *TMPRSS2, CTSL, CTSB* within different cell types in human healthy lung, liver, skin, intestine and kidney tissues. For each identified cell types in the 5 human healthy tissues analysed, the expression of the SARS-CoV2 receptor *ACE2,* the proposed alternative SARS-CoV-2 receptor CD147 (*BSG*), the serine protease TMPRSS2 and two alternative proteases involved in SARS-CoV-2 entry process were summarized in this table. -: no expression, -/+ : very few cells expressed, +: low expression, ++: high expression, +++ : very high expression.

| Tissue Types | Cell Types | *ACE2* | *BSG* | *TMPRSS2* | *CTSL* | *CTSB* |
| --- | --- | --- | --- | --- | --- | --- |
| Lung | Stromal | - | +++ | - | ++ | + |
|  | Immune | - | ++ | - | ++ | +++ |
|  | Epithelial | - | + | ++ | - | + |
|  | Endothelial | - | ++ | - | + | + |
| Liver | Pericyte & Myofibroblast | - | -/+ | - | - | - |
|  | Other | - | - | - | - | + |
|  | Epithelial | - | - | ++ | + | ++ |
|  | Hepatocyte | - | - | + | + | ++ |
|  | Endothelial | - | -/+ | - | +++ | ++ |
|  | Immune | - | - | - | - | ++ |
| Skin | Blood | - | - | - | - | - |
|  | Immune | - | + | - | - | ++ |
|  | Pericyte | - | ++ | - | + | + |
|  | Endothelial | - | ++ | - | + | + |
|  | Epithelial | - | + | -/+ | - | + |
|  | Fibroblast | - | ++ | - | ++ | ++ |
| Intestine | Transit amplifying (TA) | - | ++ | + | - | + |
|  | Stem Cell | - | + | + | -/+ | + |
|  | Progenitor | - | ++ | + | - | + |
|  | Paneth-like | - | +++ | ++ | - | + |
|  | Goblet | - | - | + | - | - |
|  | Enterocyte | ++ | ++ | ++ | - | + |
|  | Enteroendocrine | - | + | - | - | ++ |
| Kidney | Collecting duct intercalated | - | - | - | - | - |
|  | Distal tubule | - | - | - | - | - |
|  | Proximal straight tubule | + | +++ | - | + | ++ |
|  | Collecting duct principal | -/+ | ++ | - | + | ++ |
|  | Immune | - | + | - | -/+ | + |
|  | Proximal tubule | - | ++ | - | + | + |
|  | Proximal convoluted tubule | -/+ | +++ | - | ++ | +++ |
|  | Glomerular parietal epithelial | - | + | - | + | ++ |

**Figure 1.**
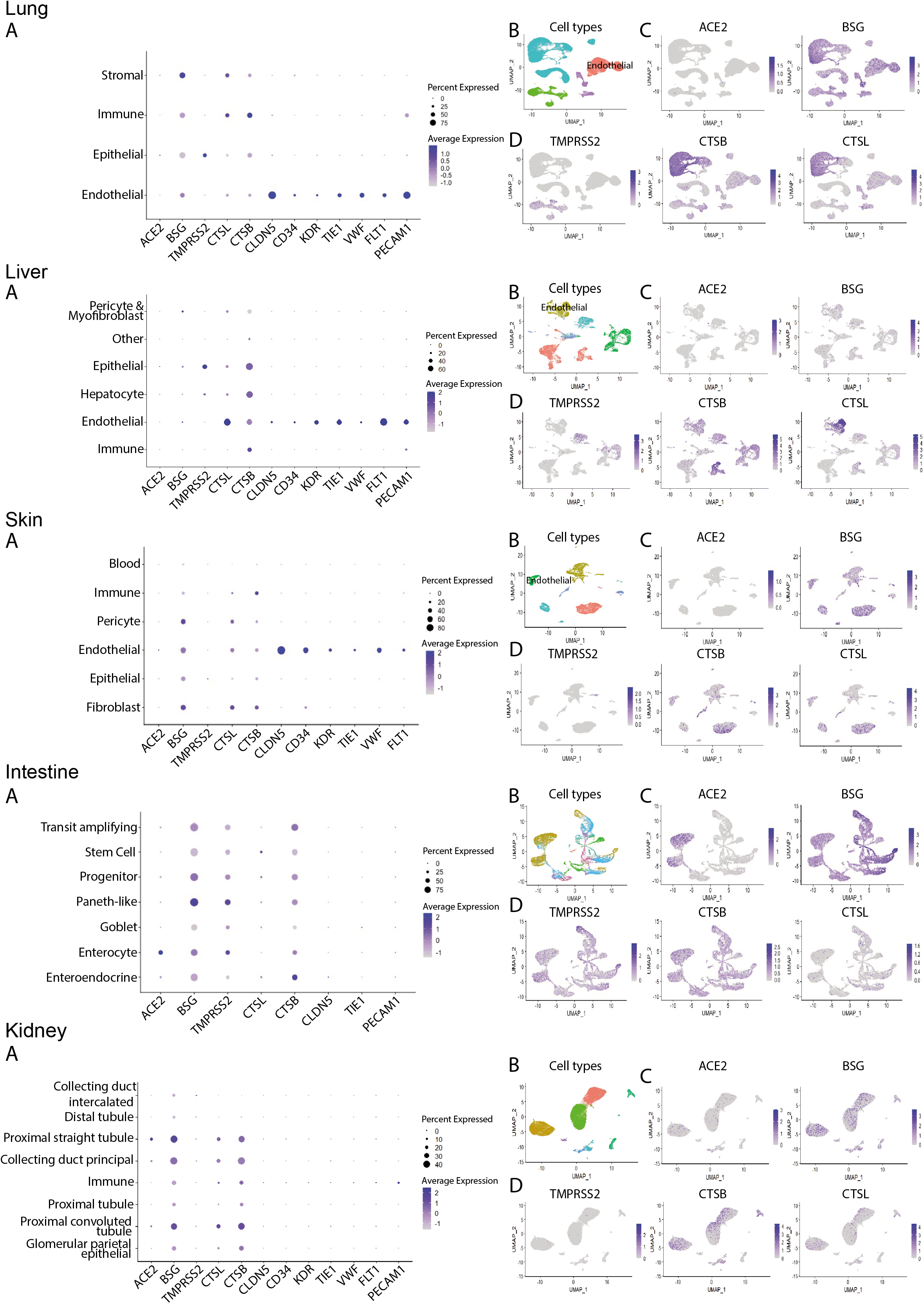
Expression of *ACE2, BSG* and *TMPRSS2, CTSL* and *CTSB* in 5 human healthy tissues. For each single-cell RNA sequencing (scRNAseq) tissue datasets, a dot plot was generated showing the expression of the SARS-CoV-2 receptor *ACE2,* the proposed alternative SARS-CoV-2 receptor CD147 (*BSG*), the serine protease TMPRSS2, two alternative proteases involved in SARS-CoV-2 entry process *CTSL* and *CTSB,* and several well known vascular endothelial markers : *PECAM1, FLT1, VWF, TIE1, KDR, CD34* and *CLDN5.* The size of the dots indicates the proportion of cells while the colour indicates log2 normalized expression of each gene (A). A UMAP plot showing the different cell types in each scRNAseq tissue dataset was generated (details in Supplementary Figure 1) and the endothelial cell populations were indicated when identified from the dataset (B). For the visualization at the single cell level, expression UMAP plots for *ACE2* and *BSG* were generated. Colour bar indicates log2 normalized expression (C). Expression UMAP plots for *TMPRSS2, CTSB* and *CTSL* were also generated. Colour bar indicates log2 normalized expression (D).

It has recently been proposed that CD147 (*BSG),* a cell surface receptor, can also facilitate entry of SARS-Cov-2 into cells (K. Wang et al. 2020) and therefore we examined expression of this transcript across the datasets. In contrast to *ACE2,* we observed that CD147 was widely expressed in these tissues including in endothelial cells in all of the datasets for which these could be identified i.e. lung, liver, and skin (Figure 1A, B, C).

## CONCLUSION

In summary, we report that CD147 but not *ACE2* is detectable in vascular endothelial cells within single cell sequencing datasets derived from multiple tissues in healthy individuals. *ACE2* expression has previously been reported by RT-PCR in the rat heart, however it was not demonstrated that this was localized to the vascular endothelium (Ferrario et al. 2005). *ACE2* expression in renal endothelium has previously been reported on the basis of immunohistochemistry (Lely et al. 2004) and in the endothelium of small and large arteries and veins in all tissues studied (Hamming et al. 2004). Both of these studies used a polyclonal rabbit anti-ACE2 antibody produced by the same manufacturer and it is possible that staining with this antibody is not specific. Interestingly, a recent report has shown that SARS-CoV-2 can infect T-lymphocytes via an *ACE2*-independent mechanism and proposed that a novel receptor might mediate uptake into T cells (X. Wang et al. 2020) and it has been shown that the level of ACE2 expression is very low in normal respiratory mucosa (Aguiar et al. 2020). In order to reconcile the absence of *ACE2* expression with the presence of viral inclusion bodies in endothelial cells of infected patients (Varga et al. 2020) and the direct infection of engineered human blood vessel organoids by COVID-19 (Monteil et al. 2020) we must conclude that either *ACE2* is not expressed in healthy tissue but is instead induced in response to SARS-Cov2, or that SARS-Cov2 enters endothelial cells via an alternative receptor such as CD147.

## Supporting information

Web appentix

