## Supplementary material for "CD147 (*BSG*) but not *ACE2* expression is detectable in vascular endothelial cells within single cell RNA sequencing datasets derived from multiple tissues in healthy individuals": Web appentix

**SUPPLEMENTARY DATA**

**Table of Contents**

1. Publicly available datasets
2. Single-cell RNA sequencing data analysis
3. Supplementary figure

1. **Publicly available datasets**

Human skin dataset

(Tabib et al. 2018)

Human kidney dataset

(Liao et al. 2020)

Human liver dataset

(Aizarani et al. 2019)

Human lung dataset

(Travaglini et al. 2020)

Human intestine dataset

(Y. Wang et al. 2020)

2. **Single-cell RNA sequencing data analysis**

The R package Seurat was used to combine linear and nonlinear dimensionality reduction algorithms for unsupervised clustering of single cells in each of the 5 publicly available datasets studied here.

Firstly, seurat objects were created with no selection and no filtration of cells to maximise detection of low or sparse ACE2 expression. The data was normalized, before feature selection and scaling. Finally, linear and non-linear dimensionality reduction and clustering was performed. Clusters and genes of interest were visualised using uniform manifold approximation and projection (UMAP) plots. Cell types were defined using metadata provided in the liver, kidney, lung and intestine datasets. In the skin dataset, cell types were determined through the expression of canonical markers. Expression levels of the markers of interest ‘ACE2’, ‘BSG’, ‘TMPRSS2’, ‘CTSL’ and ‘CTSB’ across these cell types were visualised as dotplots.

#### 3. Supplementary figure

**Supplementary Figure 1.** UMAP plots derived from multiple scRNAseq tissue datasets: Lung, Liver, Skin, Intestine and Kidney.

UMAP plots derived from 5 scRNAseq tissue datasets were generated in order to identify the different cell types in lung, liver, skin, intestine and kidney datasets.

### Supplementary Figure 1

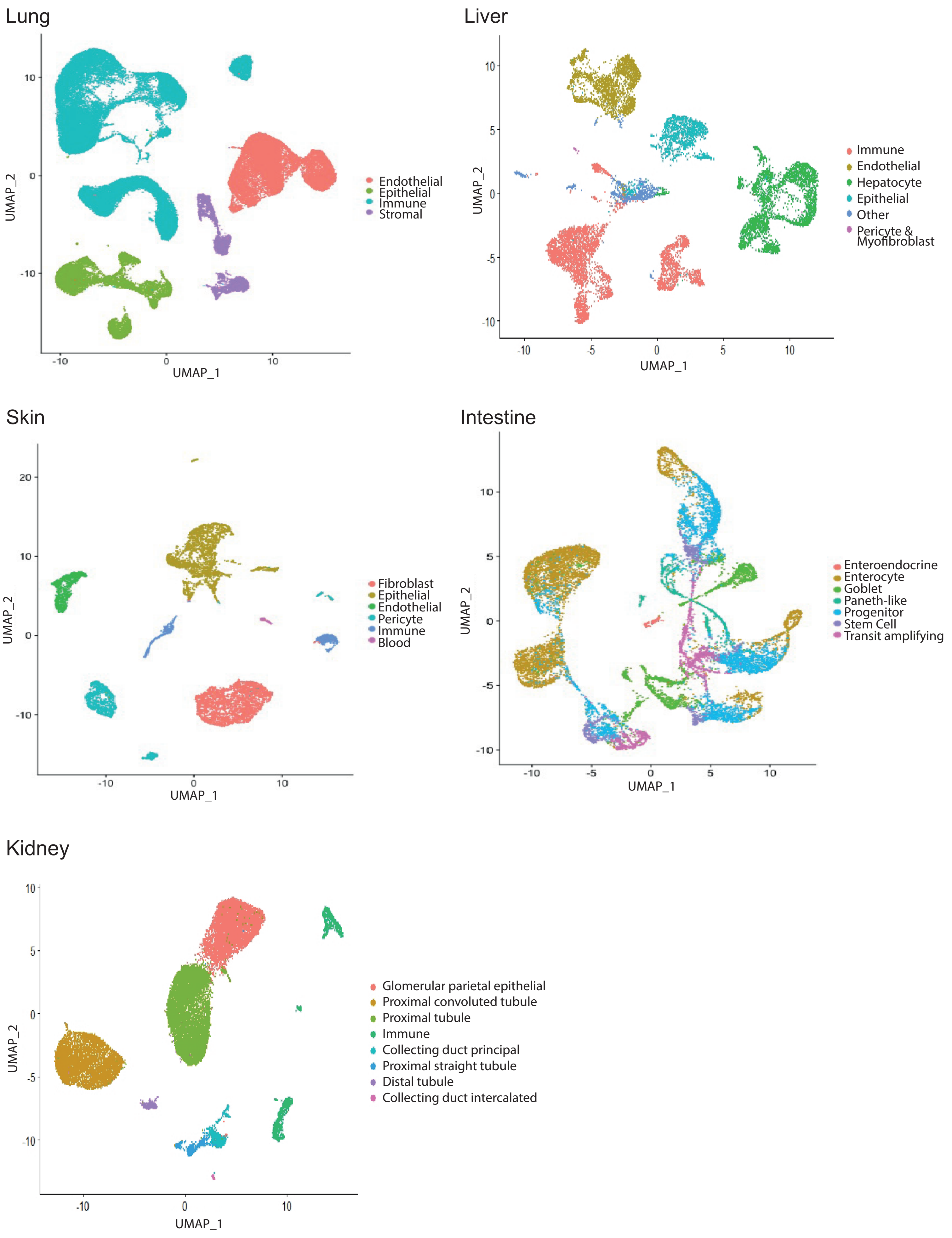
